# Bayesian *in situ* point spread function modeling enables high-fidelity three-dimensional nanoscopy in deep tissue

**DOI:** 10.64898/2026.10.27.761841

**Authors:** Jian Cao, Yumian Li, Yuyuan Qiao, Tailong Chen, Yiqing Zhang, Hao-Cheng Gao, Junyi Zhang, Lirong Qiu, Jianli Liu, Fan Xu

## Abstract

Three-dimensional single-molecule localization microscopy enables nanometer-resolution mapping of biomolecular organization in native cellular environments, but extending this capability to deep tissue requires *in situ* point spread function (PSF) modeling to account for specimen-induced optical distortions. Existing *in situ* approaches rely on a deterministic mapping between each single-molecule detection and the underlying PSF model. In deep tissue, however, specimen-induced aberrations, optical scattering and elevated background degrade the information content of single-molecule observations, posing a fundamental barrier to reliable PSF estimation. Here we introduce Bayesian *in situ* PSF modeling (BIM), a computational framework that explicitly models uncertainty in emitter axial position and propagates it through PSF reconstruction. By using an uncertainty-aware posterior fusion strategy, BIM retrieves accurate specimen-matched PSF models directly from raw blinking data under native imaging conditions. We demonstrate BIM by resolving hollow mitochondrial membranes, amyloid-β fibrils, dendrites and spines with high fidelity. When combined with adaptive optics, BIM improves high-fidelity 3D reconstruction in deep tissue at depths exceeding one hundred micrometers.

## Introduction

Elucidating the intricate molecular mechanisms of life requires visualization of biomolecular organization and interactions at nanometer resolution within their native cellular environment^1,2^. Single-molecule localization microscopy (SMLM) has revolutionized optical imaging by overcoming the diffraction limit through stochastic separation of overlapping molecules and nanometer-precision localization, enabling direct observation of biological structures at the molecular scale with spatial resolution down to ~10 nm^3–5^. However, many biological processes are governed by the three-dimensional molecular architecture of intact tissues and cannot be fully understood in isolated systems^6–8^. Three-dimensional SMLM (3D SMLM) is therefore essential for nanoscale imaging of molecular organization *in situ*, but remains challenged by the optical complexity of native tissues^9,10^.

The core principle of 3D SMLM lies in encoding and decoding the axial position of emitters through their point spread function (PSF). Specifically, axial information is encoded in an axially varying, non-degenerate 3D emission pattern, typically engineered using specific optical elements such as cylindrical lenses^11^, phase masks^12^ or multifocal planes^13^. Decoding this information requires an accurate PSF model that links the recorded emission pattern to the emitter position^14^. Early methods commonly relied on Gaussian fitting for computational efficiency, but the underlying idealized, symmetric PSF model failed to capture system-induced aberrations arising from residual optical imperfections and refractive-index mismatch^15^. To address this limitation, both analytical and numerical approaches have used pre-calibrated system models derived from measured emission patterns of fiducial markers, as well as related calibration strategies based on beads embedded in gel layers or fluorescent molecules coated on latex microspheres to characterize aberrations caused by refractive-index mismatch between immersion oil and water-based imaging media^16–20^. However, specimen-induced aberrations, optical scattering and elevated background encountered in native tissues make such models inaccurate, resulting in compromised localization precision and reconstruction fidelity^21^.

*In situ* PSF modeling is essential for accurate 3D SMLM in complex specimens, where specimen-dependent distortions compromise the accuracy of conventional PSF models^22^. Conceptually, this task can be formulated as an inverse problem with incomplete observations: although emitter images are measured, the axial positions that determine their PSF shapes remain unknown^23,24^. Inspired by the mathematical principles underlying k-means^25^ and expectation-maximization^26^, *in situ* PSF retrieval (INSPR)^23^ iteratively reconstructed the *in situ* 3D response of single emitters directly from blinking datasets, enabling high-fidelity imaging in whole-cell and tissue specimens. Furthermore, universal inverse modeling of PSF (uiPSF) ^27^ formulated PSF construction as an inverse problem and optimized PSF model parameters within an automatic differentiation framework, whereas vectorial *in situ* PSF retrieval (VISPR)^28^ combined a vectorial PSF model with maximum-likelihood phase retrieval to improve robustness to background noise. Building on these advances, *in situ* PSF modeling has been incorporated into 4Pi-SMLM to enable near-isotropic nanometer-scale localization, mapping of neuronal microcircuits in thick brain sections, and quantification of dendritic spine remodeling in the visual cortex of *Fmr1* knockout mice^29,30^. However, current *in situ* methods rely on a deterministic mapping between each detection and the underlying PSF model. In deep tissue, low photon counts and elevated background reduce the information linking PSF shape to emitter position, making the axial information of single-molecule observations inherently uncertain. This uncertain nature renders *in situ* PSF modeling underdetermined and poses a fundamental barrier to accurate 3D localization in native tissues.

Here, we develop Bayesian *in situ* PSF modeling (BIM), a computational framework that reconstructs specimen-matched *in situ* PSF models for deep-tissue 3D SMLM. BIM addresses PSF mismatch by treating emitter axial position as a latent variable and propagating posterior uncertainty directly into *in situ* PSF modeling. This uncertainty-aware strategy enables the construction of an accurate specimen-specific *in situ* PSF model from raw single-molecule datasets, thereby capturing the effective optical response of the specimen under native deep-tissue imaging conditions. We demonstrate this capability by resolving hollow mitochondrial membranes in whole cells, amyloid-β fibrils in brain sections, and dendrites and spines in cortical tissue. When combined with adaptive optics (AO), BIM further enables high-fidelity 3D reconstruction in deep tissue.

## Results

### Principle of BIM

Accurate PSF modeling is central to 3D SMLM because axial position is encoded in the shape of each single-molecule emission pattern. In deep tissue, specimen-induced aberrations, optical scattering and elevated background degrade this encoding, such that the recorded signals are no longer well described by the PSF model used for localization^22^ (Fig. 1A). BIM is a Bayesian computational framework that directly reconstructs the *in situ* 3D PSF model from raw blinking data under complex imaging conditions (Fig. 1B; Methods). Rather than enforcing a deterministic mapping between each emitter and a single axial position, BIM treats axial position as a latent variable and infers a probability distribution over candidate positions for each molecule. Through Bayesian mapping, each detected single-molecule pattern is probabilistically linked to the underlying PSF model according to its consistency with candidate emitter positions (Fig. 2A). These probabilities are then integrated through Bayesian-driven data fusion to generate empirical PSFs for comparison with model-predicted PSFs during iterative reconstruction. By representing emitter position probabilistically throughout reconstruction, BIM explicitly accounts for uncertainty in single-molecule observations and improves the robustness of *in situ* PSF modeling in deep tissue.

**Fig. 1.**
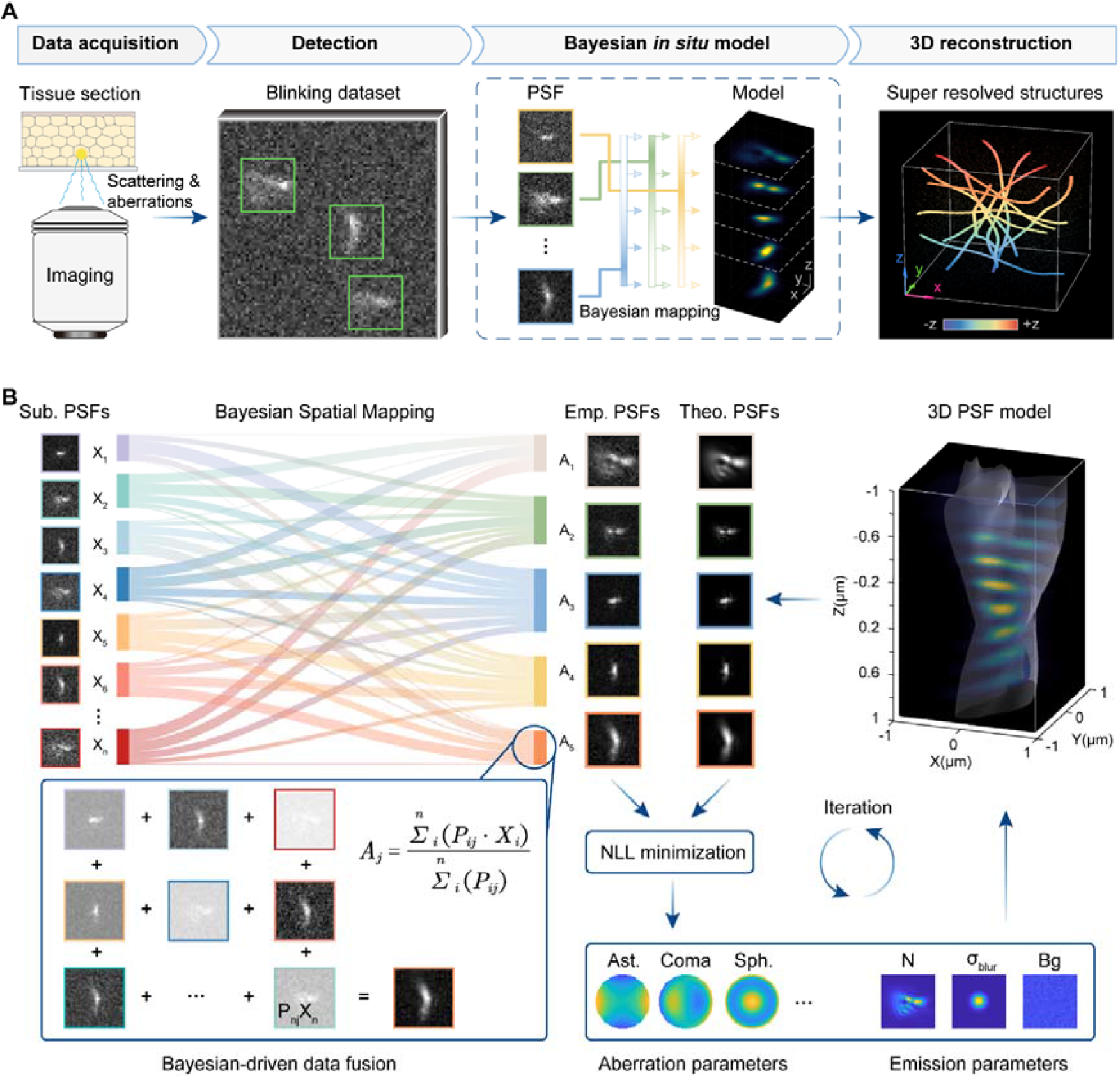
Concept and workflow of BIM. **(A)** Schematic of the overall workflow, from data acquisition and single-molecule detection to Bayesian *in situ* modeling and 3D reconstruction. **(B)** Expanded view of the Bayesian *in situ* model shown in **(A)**. Detected single-molecule patterns are probabilistically mapped to candidate axial planes, allowing each observation to contribute to multiple axial positions according to its posterior probability. These probabilities are then used for Bayesian-driven data fusion to generate high SNR empirical PSFs, with contrast indicating the contribution probability of each molecule (**Methods, Supplementary Note 1.2**). The empirical PSFs are compared with model-predicted theoretical PSFs, and maximum-likelihood optimization is used to update the Zernike coefficients of the pupil phase together with plane-wise signal intensity, background, and Gaussian blur parameters (**Supplementary Note 1.3**). The updated model is fed back into the next iteration until convergence (**Supplementary Note 1.4**). Sub. PSFs, sub-region PSFs; Emp. PSFs, empirical PSFs; Theo. PSFs, theoretical PSFs; NLL minimization, negative log-likelihood minimization; Ast., astigmatism; Sph., spherical aberration; N, photon count; Bg, background level; σ_blur_, Gaussian blur parameter.

**Fig. 2.**
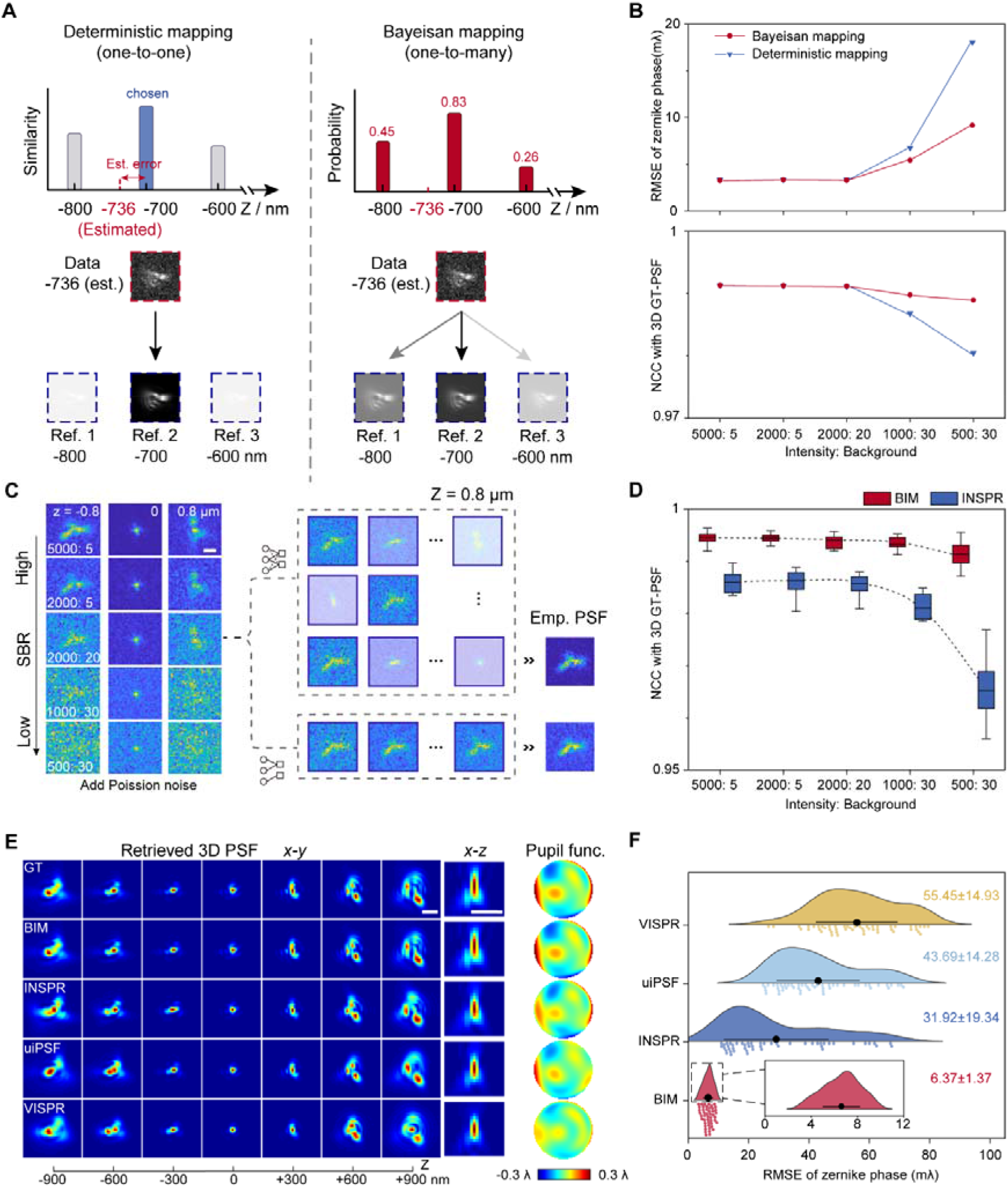
Performance quantification of BIM. **(A)** Schematic comparison of single-reference deterministic mapping and multi-reference Bayesian mapping. Deterministic mapping assigns each emitter to a single best-matched axial reference, whereas Bayesian mapping assigns posterior probabilities across candidate axial references to account for axial uncertainty. est., estimated position; Det. Map., deterministic mapping; Bay. Map., Bayesian mapping; Ref., reference axial position. **(B)** Quantitative comparison of BIM performance using Bayesian or deterministic emitter-to-axial-plane mapping under different simulation conditions. Top, root-mean-square error (RMSE) of the recovered Zernike phase. Bottom, normalized cross-correlation (NCC) between the reconstructed PSF and the ground-truth (GT) 3D PSF. **(C)** Representative simulated single-molecule patterns at decreasing signal-to-background ratio (SBR). Right, schematic comparison of Bayesian mapping- and deterministic mapping-based data-fusion strategies for generating empirical PSFs (I:bg = 1000:30, z = 800 nm). Scale bar, 1 µm. **(D)** NCC between reconstructed and GT 3D PSFs for BIM and INSPR across SBR conditions. **(E)** Representative simulation with complex aberrations. Recovered 3D PSFs for GT, BIM, INSPR, uiPSF and VISPR are shown in *x-y* and *x-z* views, together with the corresponding phase distributions. Scale bars, 1 µm. **(F)** RMSE distributions of recovered Zernike amplitudes relative to GT across 50 independent simulations (SBR = 1000:30).

BIM uses an empirical Bayesian inference framework to infer the probability distribution over candidate axial positions for each molecule^31^ (Methods). Formally, for the single-molecule emission pattern *x*_*n*_ of the n-th emitter, BIM computes the posterior probability that the emitter is located at candidate axial position *z*_*k*_, conditioned on the current forward model parameters *θ* of the pupil function.

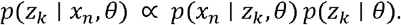

Here, *p*(*x*_*n*_ | *z*_*k*_, *θ*) represents the data-driven likelihood distribution, which quantifies how well the observed single-molecule emission pattern is explained by the PSF predicted at candidate axial position *z*_*k*_ (Supplementary Fig. 1B). In practice, this term is evaluated by fitting each detected molecule to theoretical PSF templates at discrete axial positions using maximum-likelihood estimation, yielding a likelihood profile over candidate positions. The term *p*(*z*_*k*_ | *θ*) represents the adaptive and physics-informed prior distribution, which captures axial discriminability under the current imaging conditions. Rather than assuming a uniform distribution, BIM calibrates this prior factor using inter-plane PSF similarity and Monte Carlo classification statistics under the current pupil-function model (Supplementary Fig. 1A). The resulting prior is typically narrower near focus, where the PSF carries more precise axial information, and broader at larger defocus, where reduced signal-to-noise ratio and diffraction-induced blurring diminish axial specificity (Supplementary Fig. 3A). Finally, the posterior distribution *p*(*z*_*k*_ | *x*_*n*_, *θ*) integrates the image-based evidence captured by the likelihood distribution with the optical constraints encoded in the prior (Supplementary Fig. 1C).

Bayesian-driven data fusion uses each emitter’s posterior distribution to weight the contribution of its laterally aligned emission pattern to each candidate axial plane. Weighted averaging reduces random noise and improves the signal-to-noise ratio of the empirical PSFs. Under low-SNR conditions, where axial assignment becomes less certain, retaining contributions across multiple axial planes reduces reliance on a single potentially erroneous assignment and allows axial uncertainty to be incorporated into the reconstructed 3D PSF stack (Fig. 2C).

This empirical 3D PSF stack is then used to update the pupil function through maximum-likelihood estimation phase retrieval (MLEPR)^32^ (Supplementary Fig. 4A; Supplementary Note 1.3). The pupil function is parameterized using Zernike polynomials, and the corresponding coefficients and other model parameters are optimized by minimizing a Poisson negative log-likelihood objective between the model-generated PSFs and the empirical stack. The updated pupil function generates new PSF templates for posterior estimation and weighted fusion in the next iteration. This process iterates until the retrieved model converges.

### Performance of BIM

We first assessed the core Bayesian mapping module against the conventional deterministic mapping module. Simulations were performed under aberration conditions generated from the first 21 Zernike modes in Wyant order, spanning vertical astigmatism to tertiary spherical aberration, across a range of signal-to-background ratio (SBR) conditions (Supplementary Note 3.1). The two modules showed comparable performance under high SBR, but the Bayesian mapping module yielded lower root-mean-square error (RMSE) of the estimated Zernike coefficients and higher normalized cross-correlation (NCC) for the reconstructed 3D PSFs as the SBR decreased (Fig. 2A, B).

To determine whether this improvement translated into more accurate *in situ* PSF retrieval, we compared BIM with INSPR under matched noise conditions and found that BIM better preserved PSF morphology and more faithfully reproduced the ground truth (GT) 3D PSF, with the advantage becoming more pronounced at low SBR (Fig. 2C, D; Supplementary Fig. 3B). We then evaluated the performance of BIM under simulated unknown complex aberrations with randomly sampled coefficients from the same 21 Zernike modes at a fixed SBR of 1000:30. Under these conditions, BIM achieved the lowest Zernike coefficient RMSE (6.37 ± 1.37 mλ), compared with INSPR (31.92 ± 19.34 mλ), uiPSF (43.69 ± 14.28 mλ) and VISPR (55.45 ± 14.93 mλ), and also showed closer agreement with the GT in the retrieved pupil functions, the corresponding pupil-generated 3D PSFs, the Zernike coefficients obtained from pupil decomposition, and the axial assignment accuracy (Fig. 2E, F; Supplementary Fig. 3C, D). Robust lateral registration of individual single-molecule images before posterior-weighted fusion is also important, particularly when the current PSF model does not fully match the observed emission patterns (Supplementary Fig. 2A). We therefore compared BIM’s likelihood-based fitting with NCC registration under matched and mismatched PSF models. BIM yielded lower lateral registration RMSE, with a more pronounced advantage under model mismatch (Supplementary Fig. 2B, C).

Furthermore, to mimic aberrations in thick-sample imaging, we introduced controllable wavefront distortions by placing a deformable mirror (DM) in the pupil plane. The resulting bead images were used to evaluate the fidelity of the BIM forward model, in which MLEPR serves as the computational module for modeling the PSF in a continuous Zernike space. Compared with conventional phase-retrieval methods, this BIM forward model more accurately recovered DM-induced aberrations, particularly for high-order modes (Supplementary Fig. 4A). In biplane SMLM simulations under refractive-index mismatch^17,20^, BIM showed closer agreement with the GT than INSPR in the retrieved pupil phase, the corresponding 3D PSF and the decomposed Zernike coefficients (Supplementary Fig. 4B). In simulations involving predefined, depth-dependent aberrations ranging from 0 to 72 µm^9^, BIM exhibited smaller deviations of the recovered Zernike coefficients from the GT across depths, whereas INSPR errors increased with depth (Supplementary Fig. 5A). Consistent with this result, in reconstructions of hollow elliptical structures, a static *in vitro* PSF calibrated at 0 µm produced substantial distortions at depth. INSPR reduced these artifacts, but still showed axial compression (Supplementary Fig. 5B). At 72 µm, the full width at half maximum was 10.7 nm for BIM and 10.5 nm for the GT, compared with 24.5 nm for INSPR. Peak-to-peak distance analysis showed that INSPR increasingly underestimated axial separation with depth (Supplementary Fig. 5C-E).

### 3D super-resolution imaging of thick specimens with BIM

BIM enables high-fidelity 3D super-resolution imaging in optically complex specimens by constructing specimen-matched *in situ* PSF models directly from single-molecule data acquired within the sample. We evaluated this capability in representative thick-specimen experiments spanning whole cells and brain tissue, including TOM20-labeled mitochondrial networks in COS-7 cells under DM-induced complex aberrations (Fig. 3), amyloid-β fibrils in mouse-brain sections under AO-assisted imaging (Fig. 4; Supplementary Fig. 7), and ChR2-EYFP-labeled dendrites and spines in visual cortical tissue at increasing imaging depths (Fig. 5; Supplementary Figs. 9 and 10). In each optical section, BIM derived an *in situ* PSF model tailored to the effective imaging conditions within the corresponding sample volume and incorporated this model into maximum-likelihood localization to reconstruct single-molecule emission events in 3D with high precision and reduced bias.

### BIM enables high-fidelity 3D super-resolution imaging of whole cells under complex aberration conditions

We evaluated BIM in biological specimens using an astigmatism-based 3D SMLM setup by imaging immunofluorescence-labeled TOM20 in COS-7 cells. To evaluate performance under controlled aberration conditions, we inserted a DM into the detection path and applied high-order DM modes to introduce known spatially complex aberrations that mimic severe wavefront distortions encountered in deep imaging (Fig. 3A). We then reconstructed the same raw dataset using BIM, representative *in situ* PSF methods (INSPR^23^, uiPSF^27^ and VISPR^28^), and a conventional *in vitro* calibration for direct comparison. BIM reconstructed the mitochondrial network with continuous axial organization and clear tubular connectivity across the field of view (Fig. 3B). In representative axial views, BIM resolved the hollow morphology, consistent with TOM20 labeling of the outer mitochondrial membranes^33^ (Fig. 3C-F). By contrast, reconstructions obtained with INSPR, uiPSF, VISPR and the *in vitro* calibration showed varying degrees of axial compression, profile broadening and structural distortion, reducing the visibility of the hollow geometry. Axial intensity profiles across representative mitochondrial cross-sections showed the two TOM20-positive membrane boundaries were resolved as two peaks separated by 461 nm, with fitted axial widths of 42 and 33 nm, whereas the other methods generally produced broader or less distinctly separated profiles (Fig. 3E). The pupil estimated by BIM more closely matched the DM-imposed aberration pattern while additionally accounting for sample-dependent distortions (Fig. 3G; Supplementary Fig. 6), suggesting that improved model fidelity contributed to the more accurate reconstruction.

**Fig. 3.**
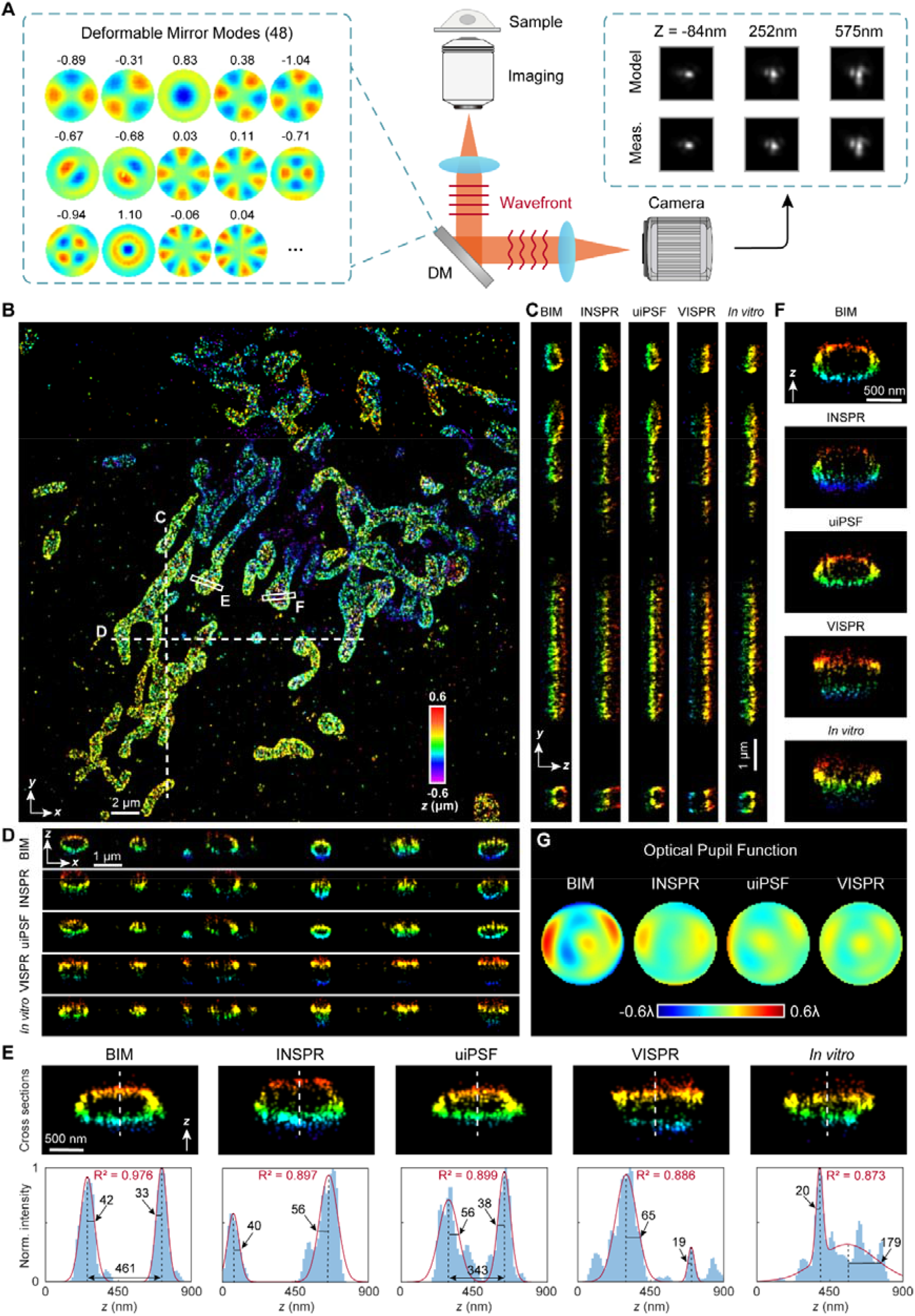
3D super-resolution imaging of TOM20-labeled mitochondria under deformable mirror (DM)-induced complex aberrations. **(A)** Experimental scheme. A DM was used to introduce high-order aberrations in an astigmatism-based 3D SMLM setup. Representative PSFs generated from the BIM-retrieved model and the corresponding measured PSFs are shown at three axial positions. DM, deformable mirror; Meas, measurement. **(B)** Overview of TOM20-labeled mitochondrial networks in COS-7 cells at the coverslip, imaged under DM-imposed complex aberrations and reconstructed with BIM. Localizations are color-coded by axial position. Scale bars, 2 µm. **(C, D)** *y-z* and *x-z* views of the selected regions indicated by white dashed lines in **(B)**, reconstructed using BIM, INSPR, uiPSF, VISPR and an *in vitro* astigmatic model (1.2 λ/2π). Scale bars, 1 µm. **(E, F)** Vertical cross-sections of selected positions (white rectangles; slice width, 200 nm) in **(B)**, and corresponding axial intensity profiles. Scale bars, 500 nm. Norm., normalized. R^2^, goodness of fit. **(G)** Retrieved phase distributions using different *in situ* methods (recovered 3D PSFs are shown in **Supplementary Fig. 6**).

### BIM resolves amyloid-β **fibrils in mouse brain sections with adaptive optics**

The three-dimensional organization of amyloid-β (Aβ) fibrils is important for understanding plaque architecture in Alzheimer’s disease^34,35^. In brain tissue, plaques are embedded within dense neuropil, coupling fibrillar ultrastructure to local tissue architecture and optical heterogeneity^36^. Imaging Aβ fibrils in brain sections at high resolution therefore provides a biologically relevant context to assess deep-tissue 3D SMLM. We evaluated BIM by imaging immunofluorescence-labeled Aβ fibrils in 125-µm-thick brain sections from 5XFAD mice, a transgenic Alzheimer’s disease model with prominent amyloid plaque pathology, under AO-assisted imaging conditions^6^ (Fig. 4; Supplementary Fig. 7). AO was used during acquisition to optimize single-molecule emission patterns, reducing wavefront distortion and improving the axial evolution of single-molecule PSFs (Fig. 4A, B). BIM reconstructed dense fibrillar networks with continuous morphology across the imaging volume (Fig. 4C). In representative *x-y* regions, BIM more clearly resolved adjacent fibrils and local separations between neighboring strands, whereas INSPR showed increased broadening or partial merging of nearby structures (Fig. 4D, E). The corresponding intensity profiles across these features showed that BIM resolved adjacent strands with fitted peak widths of 25 and 25 nm without AO correction and 15 and 24 nm after AO correction, compared with 24 and 42 nm after AO correction for INSPR. Consistent with these observations, *x-z* visualizations further showed that BIM preserved two axially separated fibrillar features with fitted peak widths of 51 and 52 nm, whereas INSPR produced asymmetric broadening, with corresponding widths of 95 and 61 nm (Fig. 4F). Across measured fibrils, BIM yielded lateral and axial fibril width values of 41 ± 9 nm and 75 ± 12 nm, compared with 59 ± 10 nm and 103 ± 19 nm for INSPR (Fig. 4G, H). For the AO-assisted Aβ reconstruction, Fourier shell correlation (FSC) analysis yielded a resolution of 68 nm for BIM, compared with 80 nm for INSPR (Fig. 4I; Supplementary Fig. 8). Additional Aβ volumes acquired under the same AO-assisted conditions showed similar trends (Supplementary Fig. 7). Together, these results indicate that BIM improves reconstruction fidelity for Aβ fibrils in aberration-prone brain tissue under AO-assisted imaging.

**Fig. 4.**
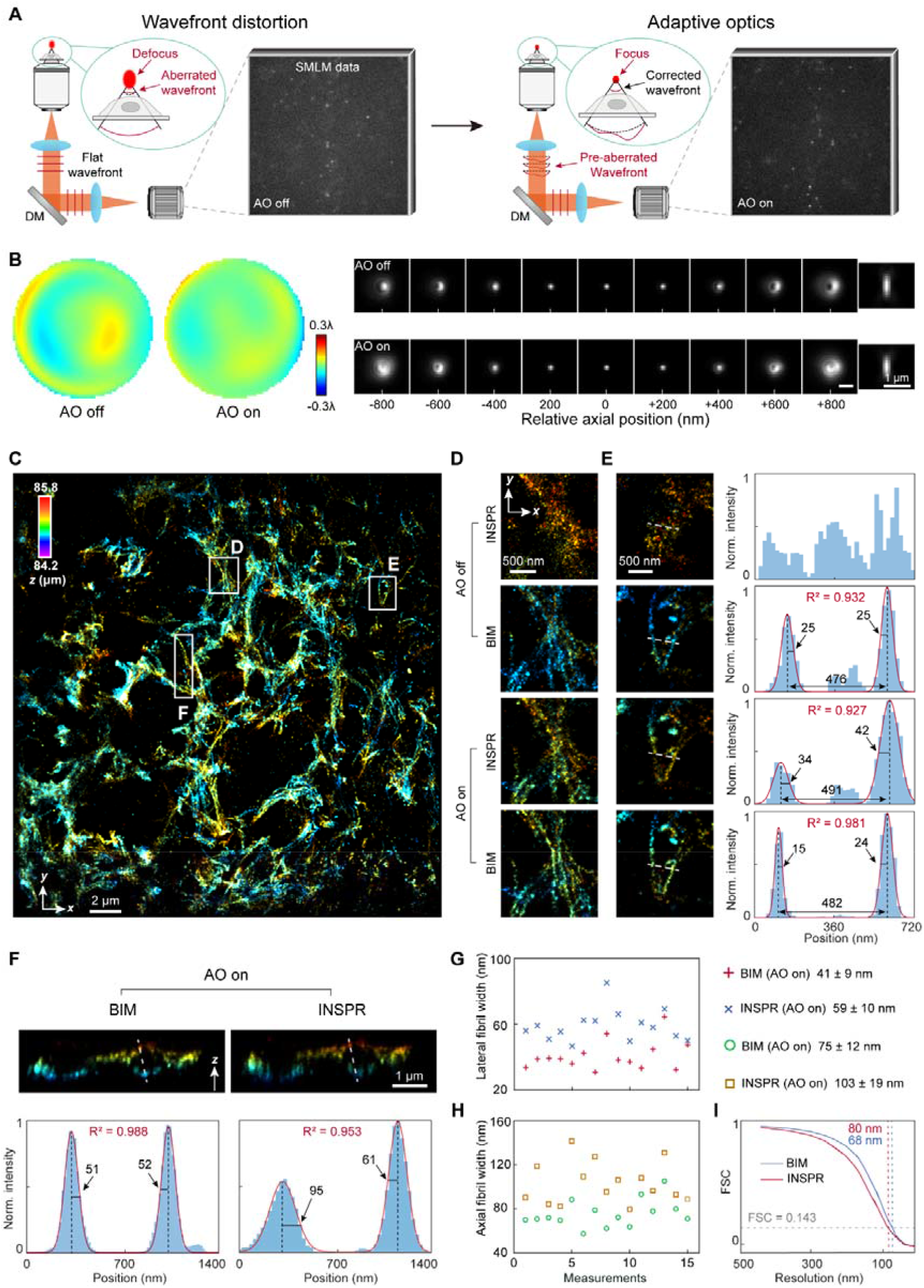
3D super-resolution imaging of immunofluorescence-labeled amyloid-β fibrils in 125-µm brain sections from 7.5-month-old 5XFAD mice. **(A)** Schematic of adaptive optics (AO) in SMLM, in which sample-induced wavefront distortions were estimated and compensated using a deformable mirror (DM). **(B)** Retrieved phase distributions and representative single-molecule PSFs before and after AO correction across the axial range. Scale bars, 1 µm. **(C)** Amyloid-β fibril in mouse brain sections imaged by SMLM with AO and reconstructed with BIM at a depth of 85 µm from the coverslip surface. Localizations are color-coded by axial position. Scale bar, 2 µm. **(D, E)** Enlarged *x-y* views of the selected regions indicated by white dashed rectangles in **(C)**, comparing reconstructions obtained without and with AO using BIM and INSPR; the corresponding intensity profiles are shown on the right. Scale bars, 500 nm. **(F)** Representative *x-z* view of the selected regions indicated by white dashed rectangles in **(C)**, comparing reconstructions obtained with AO using BIM and INSPR; the corresponding axial intensity profiles are shown below. Scale bar, 1 µm. **(G, H)** Comparison of the distributions of lateral **(G)** and axial **(H)** fibril width measured from 15 fibrils reconstructed with AO using BIM and INSPR. **(I)** Comparison of Fourier shell correlation (FSC) resolution analysis of reconstructions using BIM and INSPR. Resolution values were determined from a 5 × 5 µm^2^ central region of interest, averaged over three independent data splits at the 1/7 threshold (full analyses are shown in **Supplementary Fig. 8**).

### BIM resolves dendrites and spines in mouse brain sections

Dendrites and their associated spines are key structural features of cortical circuits, and resolving them in intact brain tissue is important for linking nanoscale morphology to circuit function^37–39^. Their visualization remains challenging because dendritic spines are submicrometer structures and imaging depth introduces substantial aberrations^40^. To assess BIM on these structures, we imaged visual cortical tissue labeled with Channelrhodopsin-2 enhanced yellow fluorescent protein (ChR2-EYFP), a membrane-associated optogenetic fusion protein that delineates neuronal surface architecture^41^ (Fig. 5; Supplementary Figs. 9 and 10). At a depth of 11 µm, where aberrations were relatively weak, BIM reconstructed extended dendritic segments with clearly resolved spine protrusions (Supplementary Fig. 9C, D). Representative *x-z* sections revealed hollow annular membrane profiles in dendritic shafts, consistent with membrane-associated labeling, and cross-sectional analysis along the spine axis further resolved the transition from narrow necks to enlarged heads (Supplementary Fig. 9E-H). At a depth of 67 µm, BIM still resolved distinct annular membrane profiles in dendritic shafts without AO correction, enabling deep-tissue imaging beyond 60 µm within a 150-µm-thick mouse brain section (Supplementary Fig. 10B-D).

When imaging deeper into tissue, AO improved reconstruction fidelity by reducing sample-induced aberrations. Under AO-assisted imaging, BIM reconstructed continuous dendritic shafts and spine protrusions at depths of 67 µm and 126 µm (Fig. 5A, D; Supplementary Fig. 10A). In axial sections, BIM resolved the hollow tubular profile of dendritic shafts and spines with separated opposing membrane boundaries, whereas INSPR and uiPSF yielded broader axial profiles with reduced membrane separation (Fig. 5B, E). These differences were supported by axial intensity profiles, which showed sharper dual-peak separation for BIM at both depths. Individual spines resolved by BIM exhibited clear head-neck organization in three dimensions. In lateral views, narrow necks connected enlarged heads, whereas in axial sections the head region showed broader membrane-separated profiles than the adjoining neck. Cross-sectional area measurements along the spine axis were consistent with this morphology, revealing constricted necks that expanded into bulbous heads and were well described by sigmoidal fits (Fig. 5C, F and G; Supplementary Fig. 11). 3D alpha-shape modeling of the localization clouds further supported mushroom-like spine geometries at both depths, and additional dendritic volumes acquired under the same AO-assisted conditions showed similar results (Supplementary Fig. 10B-F). Together, these results show that BIM enables robust recovery and quantification of dendritic spine morphology deep in brain tissue.

**Fig. 5.**
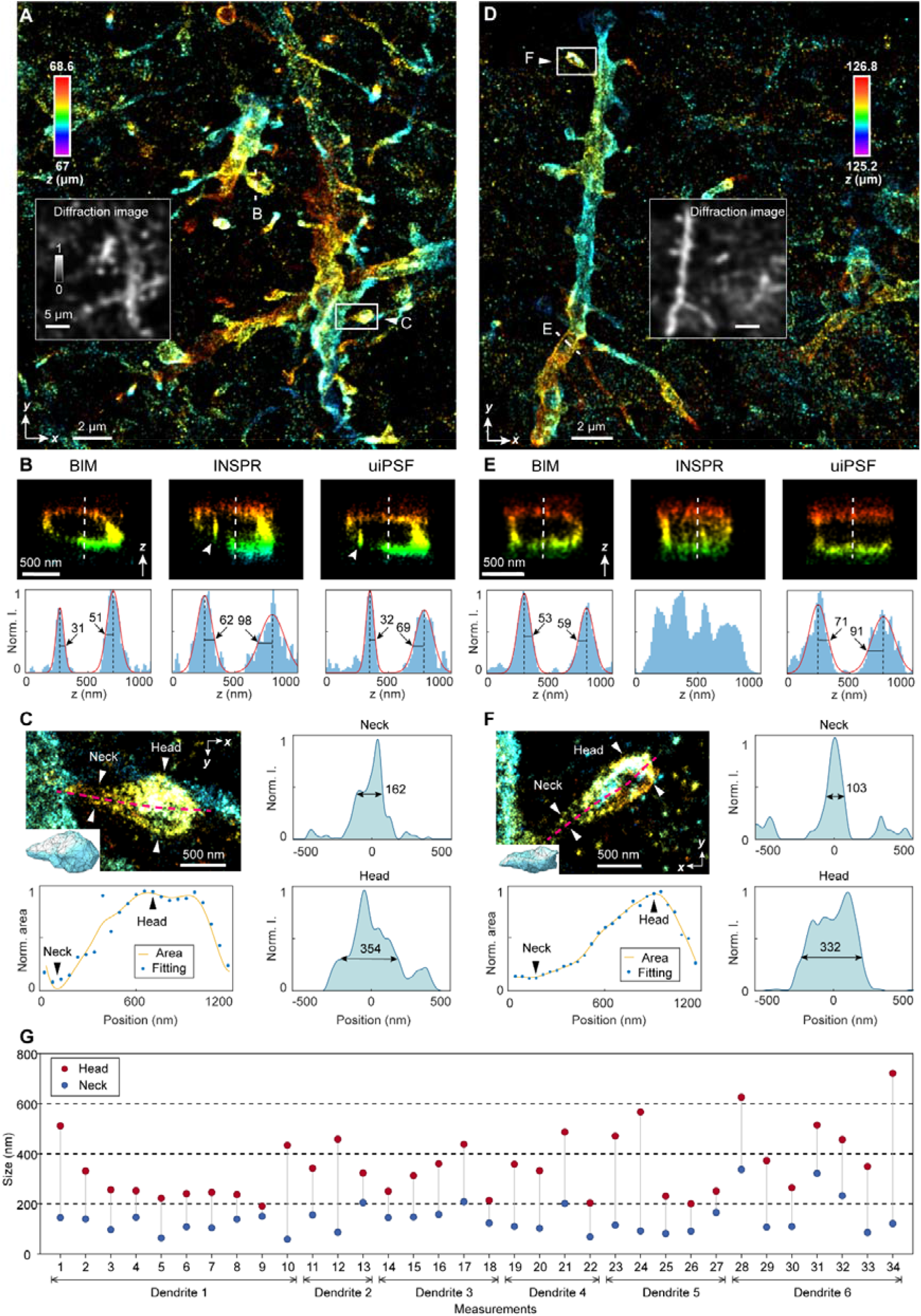
3D super-resolution imaging of dendrites and spines in immunofluorescence-labeled ChR2-EYFP in 150–250-µm-cut brain sections of 7-week-old mice. **(A, D)** Overview of 3D reconstructions of ChR2-EYFP-labeled dendrites and spines in mouse visual cortical sections imaged under AO-assisted SMLM and reconstructed with BIM at depths of 67 µm **(A)** and 126 µm **(D)** from the coverslip. Localizations are color-coded by axial position. Insets, corresponding diffraction-limited images of the same regions. Scale bars, 2 µm; inset scale bars, 5 µm. **(B, E)** Representative *x-z* views of dendritic shaft regions indicated in **(A)** and **(D)**, comparing reconstructions obtained with BIM, INSPR and uiPSF. Bottom, corresponding axial intensity profiles extracted along the dashed vertical lines. Numbers indicate the peak-to-peak separations obtained from the profiles. Arrowheads mark broadened or poorly resolved membrane features in the comparison methods. Scale bars, 500 nm. **(C, F)** Quantitative analysis of representative individual dendritic spines selected from the regions shown in **(A)** and **(D)**. Left, magnified depth-colored views with the spine centerline indicated by the dashed magenta curve; inset, corresponding 3D alpha-shape model. Bottom left, normalized cross-sectional area along the spine axis with fitted trend. Right, transverse profile measurements at the neck and head regions, with the estimated widths indicated. Scale bars, 500 nm. **(G)** Summary of measured spine-neck and spine-head sizes for 34 spines from six dendrites (representative dendritic spines are shown in **Supplementary Fig. 11**).

## Discussion

Accurate PSF modeling is central to 3D SMLM, with axial position encoded in the shape of each single-molecule emission pattern. In deep tissue, however, specimen-induced aberrations, refractive-index heterogeneity and elevated background degrade this encoding and increase mismatch between measured emission patterns and the PSF model used for localization. BIM addresses this challenge by treating axial position as a latent variable and propagating posterior uncertainty directly into *in situ* PSF reconstruction. This uncertainty-aware strategy improves the correspondence between the optical forward model and the effective imaging conditions within the specimen, particularly under optically complex conditions in which PSF mismatch is most severe. In our simulations and experiments, these gains translated into more faithful recovery of biological ultrastructure, including hollow mitochondrial membranes, amyloid-β fibrils in brain tissue, and membrane-defined dendritic shafts and spines. Combined with AO, BIM further supported high-fidelity 3D imaging in tissue at depths of up to ~126 µm. These results establish uncertainty-aware *in situ* PSF modeling as a practical framework for extending quantitative 3D SMLM to deeper and more heterogeneous tissue environments.

A central goal in biological imaging is to extend nanoscale measurements to greater depths, larger specimen volumes and richer molecular content while maintaining quantitative accuracy. Extending 3D SMLM across such imaging regimes will require accurate PSF modeling under increasingly demanding optical conditions, and BIM may provide a useful framework for these developments. Integration with tissue clearing^42^ may enable imaging at greater depths by reducing scattering and refractive-index heterogeneity. Extension to multicolor imaging^43,44^ will require treatment of wavelength-dependent PSF variation and chromatic aberration. Imaging over larger fields of view^45^ will require compensation for field-dependent PSF distortions arising from field curvature and spatially heterogeneous aberrations. Higher spatial precision can be achieved through interferometric implementations^30,46^ such as 4Pi-SMLM, which require particularly accurate modeling of interference and specimen-dependent aberrations.

In summary, BIM provides a statistically grounded and physically interpretable framework for *in situ* PSF modeling in 3D SMLM. Beyond improving specimen-matched reconstruction in optically complex tissues, we anticipate that it may facilitate the application of quantitative super-resolution imaging to more intact and heterogeneous biological specimens.

## Methods

### Overview of the BIM framework

Bayesian *in situ* point spread function modeling (BIM) reconstructs specimen-matched 3D PSF models directly from raw single-molecule blinking data. The central unknown in this inverse problem is the axial position of each detected emitter: every molecule contributes a noisy PSF image, but its z position must be inferred from the same data used to build the model. BIM therefore treats axial position as a latent variable and estimates a posterior probability over candidate axial planes. These posterior probabilities are used to fuse aligned single-molecule images into an empirical 3D PSF stack, which is then fitted with a physical pupil-function model. Iterating posterior inference and physical model refinement yields an *in situ* PSF model matched to the local imaging condition. The final model is then used for 3D localization of all detected emitters. Details of the forward model, maximum-likelihood estimation phase retrieval and iterative refinement are provided in Supplementary Note 1.

### Concept of Bayesian spatial mapping

In SMLM, the *in situ* PSF is not observed directly as a complete three-dimensional object, but is sampled sparsely through many stochastic blinking events distributed across the axial range. Each detected emitter image can be regarded as a noisy observation of the same underlying 3D PSF evaluated at an unknown axial position. Raw blinking data therefore contain the information required for *in situ* PSF recovery, but this information is only partially observed: the emission pattern of each molecule is measured, whereas its axial coordinate remains latent and must be inferred from the same image. This incomplete-observation structure makes PSF reconstruction fundamentally different from standard image averaging or calibration-based PSF estimation. Existing reconstruction strategies often assign each emitter to a single axial plane before model updating. Although this deterministic mapping simplifies the reconstruction procedure, it does not explicitly preserve axial ambiguity arising from low signal-to-noise ratio, elevated background, limited axial discrimination, or aberration-induced PSF degeneracy. To address this limitation, BIM formulates Bayesian spatial mapping as a probabilistic module that estimates, for each emitter, a posterior distribution over candidate axial planes under the current forward model and then propagates this uncertainty into empirical PSF reconstruction. In this framework, Bayesian spatial mapping serves two coupled functions within BIM: it infers the probabilistic correspondence between individual emitter images and the current axial PSF stack, and it converts these probabilistic assignments into a posterior-weighted empirical representation for subsequent model refinement.

Let *x*_*n*_ denote the observed image patch of the n-th emitter, and let *z*_*k*_, *k* = 1, …, *K*, denote the candidate axial planes in the discretized PSF stack. At iteration *t*, the current optical model *θ*^(*t*)^ generates the corresponding model-predicted PSFs from the shared pupil function *P*(*k*_*x*_, *k*_*y*_). BIM estimates the posterior probability that emitter n originated from plane *z*_*k*_,

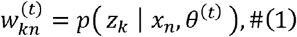

According to Bayes’ theorem,

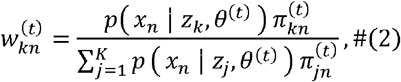

where *p*(*x*_*n*_ | *z*_*k*_, θ^(*t*)^) is the data-driven likelihood, and 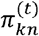 is the physics-informed axial prior factor used for emitter n at candidate plane *z*_*k*_. This formulation allows a single emitter to contribute to multiple axial planes according to posterior mass, rather than being forced into a deterministic mapping.

### Data-driven likelihood calculation

The likelihood term quantifies how well the observed image patch *x*_*n*_ agrees with the theoretical PSF predicted at candidate axial plane *z*_*k*_ under the current optical model θ^(*t*)^ (Supplementary Fig. 1B). For each emitter and each candidate plane, BIM fits the lateral displacement, photon count and background while holding the candidate axial coordinate fixed. The fitted model prediction at pixel *r* is denoted by *μ*knr, and the measured pixel value is denoted by *x*_*n,r*_. Because single-molecule fluorescence measurements are governed by photon-counting statistics, each pixel is modeled as a Poisson-distributed observation, *x*_*n,r*_ ~ Poisson(*μ*_*knr*_). Under this noise model, the likelihood can be expressed from the Poisson log-likelihood-ratio loss as.

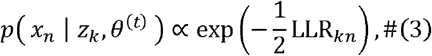

Here, LLR_*kn*_ (log-likelihood-ratio) measures the agreement between emitter *n* and the fitted model at candidate plane *z*_*k*_, and is computed as

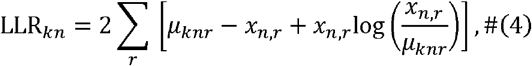

Lower LLR_*kn*_ values indicate closer agreement between the observed single-molecule emission pattern and the model-predicted PSF. Repeating this maximum-likelihood fitting across all candidate axial planes yields a likelihood vector for each emitter, and repeating it across all detected emitters yields the likelihood matrix used for Bayesian posterior estimation. This formulation assigns the likelihood scale through the Poisson noise model.

### Physics-informed prior calibration

The physics-informed axial prior factor is constructed from the current optical forward model to account for variations in axial discriminability across the imaging range (Supplementary Fig. 1A). At some axial positions, small defocus changes produce clearly distinguishable PSF morphologies, whereas at others, similar emission patterns make neighboring planes difficult to distinguish, particularly under strong aberrations or reduced signal-to-background ratios. To characterize this model-dependent ambiguity, BIM generates noise-free reference images at predefined axial planes and computes an inter-plane similarity matrix using normalized cross-correlation. In parallel, Monte Carlo simulations generate Poisson-noise realizations at each plane using signal and background levels estimated from the data. Each realization is assigned to the reference plane with the highest correlation, yielding a confusion matrix whose entries record the assignment frequencies for each generating plane. For each generating plane, a zero-intercept least-squares fit relates the valid assignment frequencies to the corresponding similarity matrix, yielding an axial-plane-specific scaling coefficient. BIM applies the calibrated exponential transformation to each similarity profile to construct the prior-factor matrix. The prior is narrow when axial discriminability is high and broader when the PSF manifold is locally degenerate (Supplementary Fig. 3A).

### Bayesian posterior computation

The posterior distribution is obtained by combining the data-driven likelihood with the calibrated prior and normalizing across candidate axial planes (Supplementary Fig. 1C). For emitter, BIM computes the posterior probability,

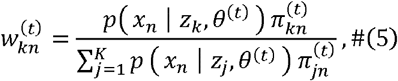

In implementation, this calculation is performed in the log domain for numerical stability by combining 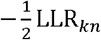 with 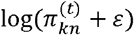, subtracting the maximum log weight for each emitter, exponentiating and normalizing over *k*. The resulting posterior matrix defines the probabilistic weights used for posterior-weighted data fusion in the BIM optimization loop (Fig. 1B; Supplementary Fig. 1C). Emitters with distinctive PSF morphology produce concentrated posterior distributions, whereas ambiguous or low-SNR emitters distribute posterior mass over neighboring planes.

### Posterior-weighted empirical PSF reconstruction

The posterior probabilities are used to reconstruct an empirical 3D PSF stack by weighted fusion of laterally aligned single-emitter observations. Let *I*_*n, r*_ denote the pixel value of emitter *n* after lateral registration to a common center. For each candidate axial plane *z*_*k*_, the posterior-fused empirical PSF intensity at pixel *r* is

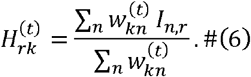

The effective probability mass 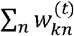 is used to determine whether plane *z*_*k*_ contains sufficient support for model refinement. Collecting 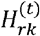 over all pixels and axial planes yields the empirical stack *H*^(*t*)^, which is passed to MLEPR. Because each emitter contributes in proportion to its posterior probability rather than through a binary label, this reconstruction preserves axial continuity and reduces sensitivity to individual classification errors.

### Extension to biplane imaging

In the biplane configuration, paired images from the two detection channels are treated as axially shifted observations of the same pupil function, with effective axial positions *z*_*k*_ ™ *δ* / 2 and *z*_*k*_ + *δ* / 2, where *δ* denotes the calibrated biplane distance. The prior calibration is performed using paired theoretical templates, and the likelihood is computed from a joint maximum-likelihood fit of the paired emitter images. Equivalently, the biplane likelihood is obtained by summing the channel-specific Poisson log-likelihoods, or by summing their corresponding LLR terms. The resulting posterior distribution 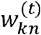 is then used to reconstruct two channel-specific empirical PSF stacks with matched posterior weights, preserving the probabilistic structure of BIM while exploiting the additional phase diversity provided by biplane detection.

## Data availability

Data underlying the results presented in this paper are not publicly available at this time but may be obtained from the authors upon reasonable request.

## Code availability

An integrated software implementation is provided to support both astigmatism-based and biplane 3D SMLM workflows. The framework includes tools for image preprocessing, Bayesian *in situ* PSF model generation, pupil-based 3D localization, drift correction, volume alignment and super-resolution rendering. Source code, documentation and example scripts required to reproduce the main analyses will be made freely available through a public GitHub repository upon publication, with further updates and additional resources provided thereafter.

## Acknowledgements

We thank Fang Huang from Purdue University for providing single-molecule datasets used in DL-AO. This work was supported by the National Natural Science Foundation of China (62272041), the National Key R&D Program of China (2023YFC3402600), the Fundamental Research Funds for the Central Universities, the Postdoctoral Fellowship Program of CPSF (GZB20250982) and the Beijing Institute of Technology Research Fund Program for Young Scholars.

## Notes

### Competing Interest Statement

The authors have declared no competing interest.

